# Efficacy of Anti-Staphylococcal Lysin, LSVT-1701, in Combination with Daptomycin in Experimental Left-Sided Infective Endocarditis Due to Methicillin-Resistant *Staphylococcus aureus* (MRSA)

**DOI:** 10.1101/2021.03.12.435219

**Authors:** David B. Huang, Eric Gaukel, Nancy Kerzee, Katyna Borroto-Esoda, Simon Lowry, Yan Q. Xiong, Wessam Abdelhady, Arnold S. Bayer

## Abstract

MRSA endovascular infections are frequently recalcitrant to treatment with standard-of-care antibiotics. Anti-staphylococcal phage lysins represent important candidate adjunctive agents against invasive MRSA infections because of both their microbicidal and anti-biofilm properties. We utilized the rabbit model of aortic valve infective endocarditis (using the prototype MRSA strain, MW2) to examine the combined efficacy of the lysin, LSVT-1701, plus daptomycin. LSVT-1701 was given at two dose-regimens (32.5 mg/kg and 50 mg/kg) with different dose-durations (single dose vs daily dose for 2 d vs daily dose for 4 d); daptomycin was given at a sub-lethal daily dose of 4 mg/kg for 4 d to maximize potential synergistic interaction outcomes. The combination of LSVT-1701 plus daptomycin was highly effective at reducing target tissue MRSA counts (cardiac vegetations, kidneys, and spleen), especially when the lysin was given for multiple days and/or at higher daily doses. Of importance, when given for four daily doses, both lysin dose-regimens in combination with daptomycin sterilized all target tissues. These findings suggest that LSVT-1701 warrants further clinical evaluation as adjunctive therapy for the treatment of invasive MRSA infections.

## Introduction

*Staphylococcus aureus* is a formidable pathogen because of its many innate virulence factors (1). Despite the availability of the current antibiotic armamentarium, *Staphylococcus aureus* infections especially those involving the endovascular system (e.g., infective endocarditis [IE], cardiac and hemodialysis device infections and others) are associated with high morbidity and mortality rates (2,3). This is particularly true when such infections are caused by multi-drug resistant strains of methicillin-resistant *S. aureus* (MRSA). For example, even MRSA strains that have a minimal inhibitory concentration (MICs) for vancomycin within the accepted Clinical Standards Laboratory Institute (CLSI) ‘susceptible’ range (i.e., ≤ 2 μg/mL) frequently fail clinical therapy with this agent (4). Thus, there is an urgent need for new therapeutic alternatives or adjuncts, which utilize novel mechanisms of action, for treating such infections.

LSVT-1701 (previously known as SAL200), is a novel, recombinantly-produced, bacteriophage-encoded lysin that specifically targets staphylococci via cell wall enzymatic hydrolysis (5). LSVT-1701 has a cell-wall-binding domain and two catalytic domains (an endopeptidase and an amidase), which degrade the staphylococcal cell wall peptidoglycan, eliciting rapid osmotic lysis, leading to an ultimate staphylocidal effect. LSVT-1701 is being developed as an adjunctive agent to combine with standard-of-care antibiotics against *S. aureus*, including both methicillin-sensitive *S. aureus* (MSSA) and MRSA. LSVT-1701 has the following important properties: **i**) it is rapidly bacteriolytic, with a narrow LSVT-1701 MIC range against a wide range of *S. aureus* isolates, including multi-resistant clinical isolates (1); **ii**) it has potent anti-biofilm activity (6); **iii**) it is synergistic with standard-of-care anti-staphylococcal antibiotics, such as vancomycin and daptomycin *in vitro* (1); **iv**) it can be dosed multiple times without dose-limiting toxicities or hypersensitivity in rodents or non-human primates, following single- and repeated-dose toxicity studies; and **v**) has a low propensity for emergence of resistance.

LSVT-1701’s anti-biofilm properties make it particularly relevant to treatment of biofilm-associated infections, such as IE and osteomyelitis. We, thus, sought to examine the *in vivo* efficacy of LSVT-1701 at multiple dose-regimens in combination with daptomycin, a standard-of-care anti-MRSA agent with proven efficacy against bacteremia and IE in humans (7). We employed a standard left-sided aortic valve IE model in rabbits, utilizing a prototypic MRSA strain, MW2, used in many prior experimental IE investigations (8,9). We examined microbiologic outcomes in distinct target tissues in this model, as well as the pharmacokinetic and pharmacodynamic drivers and target attainment values most predictive of treatment outcomes.

## Results

Baseline (pre-therapy) MRSA blood culture counts were similar in all groups, including untreated controls (between 3.18 – 3.50 log_10_ CFU/ml). All the LSVT-1701 and daptomycin combinations, as well as daptomycin alone, regimens rendered blood cultures as negative at time of sacrifice (24 h after the last dose of both study drugs).

All LSVT-1701 regimens in combination with daptomycin significantly reduced MRSA burdens in all target tissue as compared to untreated controls (**Table 1; Figures 1–3**). The reduction in MRSA counts was statistically enhanced in groups featuring both increasing LSVT-1701 doses (i.e., single doses of 50 mg/kg vs 32.5 mg/kg iv), as well as increased numbers of lysin doses (i.e., four daily doses vs a single-dose or two daily-doses) in combination with daptomycin. Of note, both the LSVT-1701 50 mg/kg and 32.5 mg/kg daily dose-strategies given for four days in combination with daptomycin sterilized all target tissues (i.e., quantitative cultures at or below the lower limit of detection of 1 log_10_ CFU/g. tissue). Also, LSVT-1701 given as a 50 mg/kg single dose, in combination with daptomycin, sterilized 4/10 vegetations, 3/10 kidney abscesses, and 7/10 splenic abscesses. In comparison, daptomycin alone or in combinations with the lower LSVT-1701 dose of 32.5 mg/kg single dose or two daily dose-regimens failed to sterilize a single target tissue sample.

**Figure 1.**
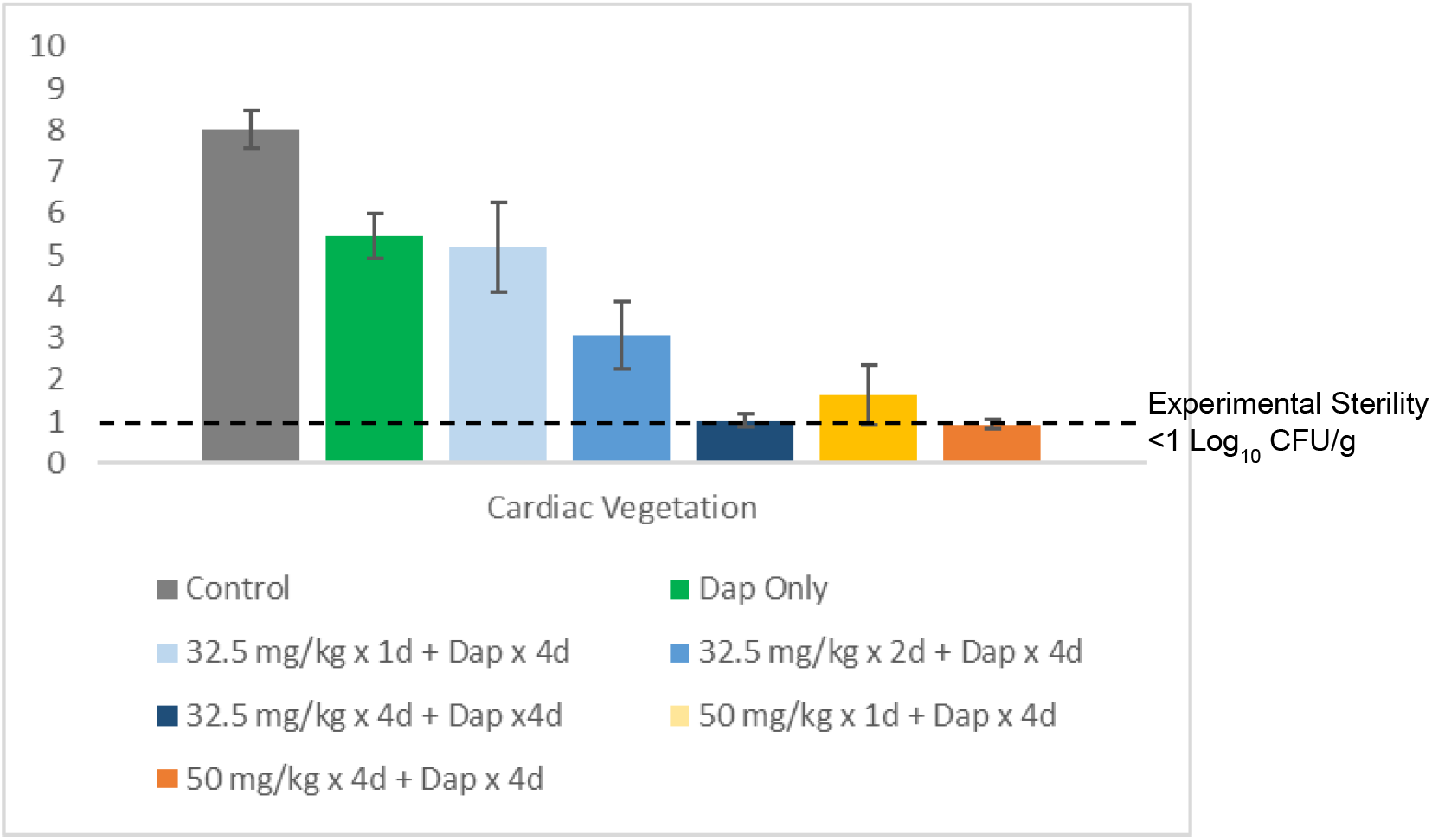
MRSA bioburdens in cardiac vegetations. Abbreviations: MRSA, methicillin-resistant *S. aureus*; Dap, daptomycin; mg, milligram; kg, kilogram; CFU, colony forming unit; g, gram. Error bars represent standard deviation.

**Figure 2.**
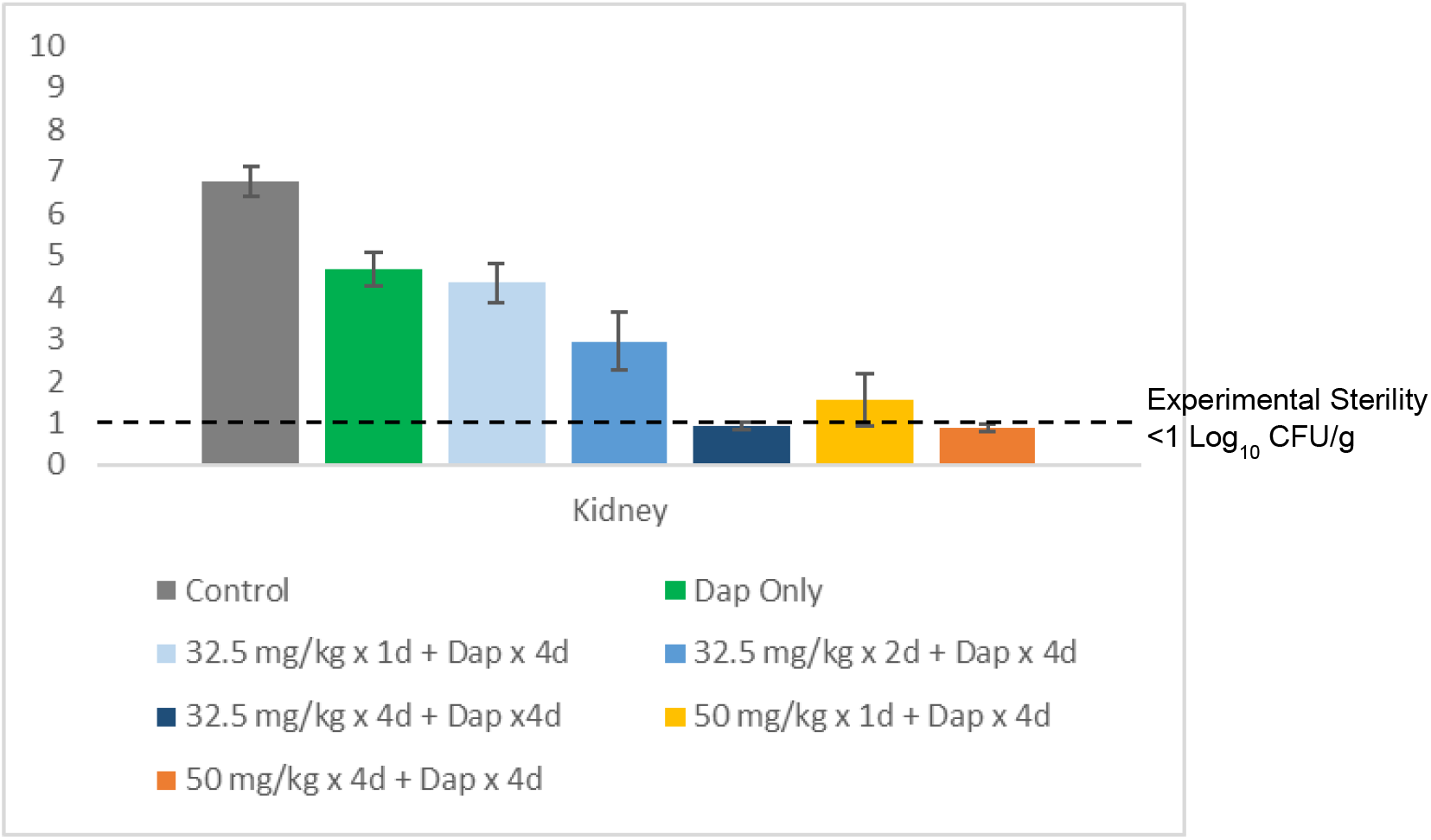
MRSA bioburdens in kidneys. Abbreviations: MRSA, methicillin-resistant *S. aureus*; Dap, daptomycin; mg, milligram; kg, kilogram; CFU, colony forming unit; g, gram. Error bars represent standard deviation.

**Figure 3.**
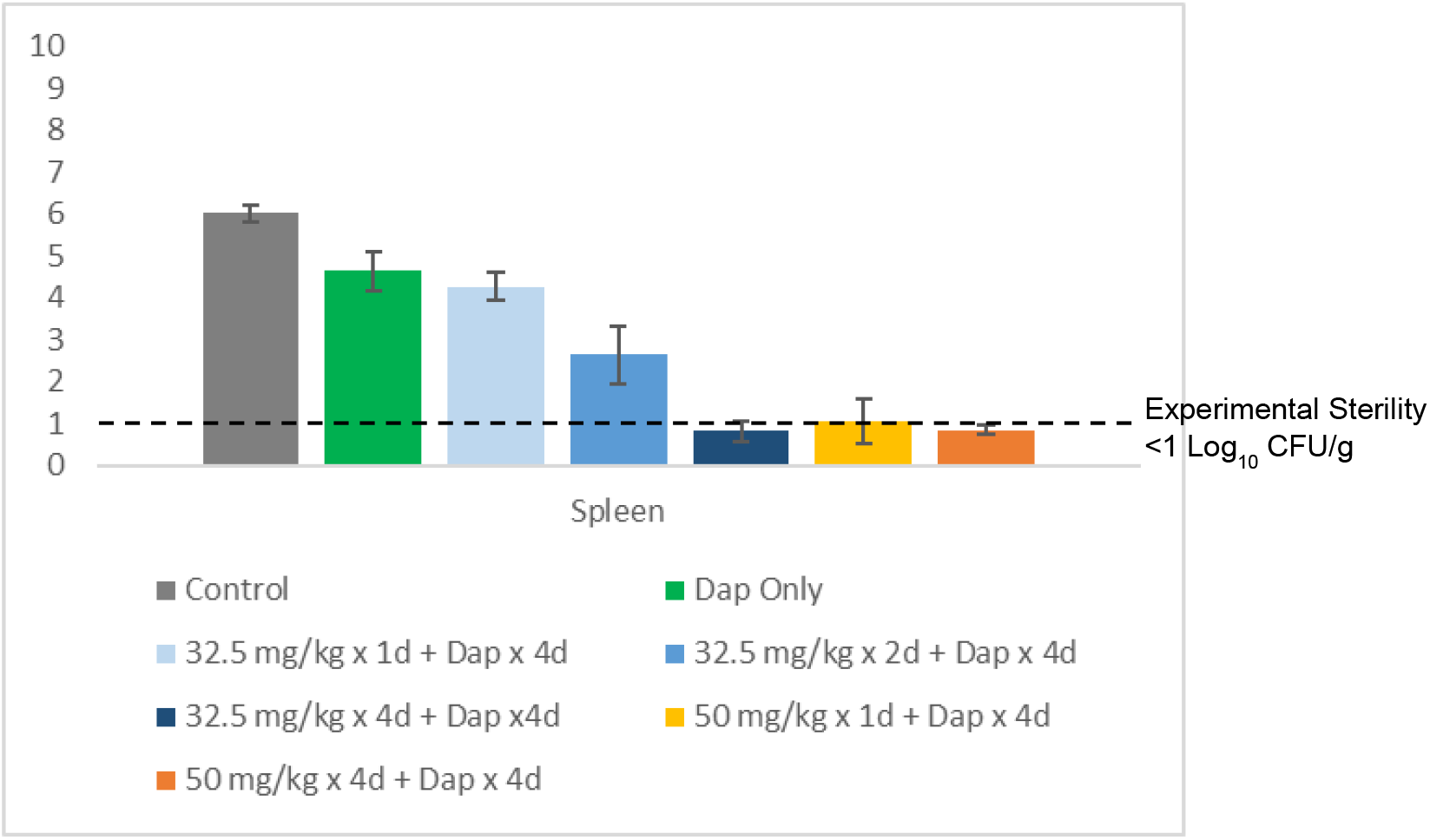
MRSA bioburdens in spleen. Abbreviations: MRSA, methicillin-resistant *S. aureus*; Dap, daptomycin; mg, milligram; kg, kilogram; CFU, colony forming unit; g, gram. Error bars represent standard deviation.

**Table 1.** MRSA bio-burdens in blood, vegetations, kidneys and spleen in rabbit IE model.

| Groups<br>(No. of animals) | Log<sub>10</sub> CFU/g. tissue (Mean ± SD)<br>(No. of sterile cultures) |  |  | Log<sub>10</sub> CFU/ml (Mean ± SD)<br>(Pre-therapy baseline) |
| --- | --- | --- | --- | --- |
|  | Vegetations | Kidneys | Spleen | Blood |
| Untreated control<br>(N=10) | 8.02 ± 0.44<br>(0/10) | 6.77 ± 0.35<br>(0/10) | 6.02 ± 0.20<br>(0/10) | 3.18 ± 0.45 |
| Dap 4 mg/kg iv QD x 4d<br>(N=10) | 5.44 ± 0.53*<br>(0/10) | 4.66 ± 0.41*<br>(0/10) | 4.64 ± 0.46*<br>(0/10) | 3.33 ± 0.40 |
| LSVT-1701 32.5 mg/kg iv single dose<br>+ Dap 4 mg/kg iv QD x 4d<br>(N=10) | 5.17 ± 1.09*,%<br>(0/10) | 4.34 ± 0.45*,%<br>(0/10) | 4.27 ± 0.33*,%<br>(0/10) | 3.22 ± 0.49 |
| LSVT-1701 32.5 mg/kg iv QD x 2d<br>+ Dap 4 mg/kg iv QD x 4d<br>(N=10) | 3.06 ± 0.81*,@,#,%<br>(0/10) | 2.95 ± 0.70*,@,#,%<br>(0/10) | 2.64 ± 0.67*,@,#,%<br>(0/10) | 3.37 ± 0.27 |
| LSVT-1701 32.5 mg/kg iv QD x 4d<br>+ Dap 4 mg/kg iv QD x 4d<br>(N=10) | 1.01 ± 0.16*,@,#<br>(10/10) | 0.91 ± 0.09*,@,#<br>(10/10) | 0.82 ± 0.23*,@,#<br>(10/10) | 3.30 ± 0.22 |
| LSVT-1701 50 mg/kg iv single dose<br>+ Dap 4 mg/kg iv QD x 4d<br>(N=10) | 1.64 ± 0.73*,@<br>(4/10) | 1.56 ± 0.61*,@,\$<br>(3/10) | 1.06 ± 0.53*,@<br>(7/10) | 3.50 ± 0.39 |
| LSVT-1701 50 mg/kg iv QD x 4d<br>+ Dap 4 mg/kg iv QD x 4d<br>(N=10) | 0.93 ± 0.10*,@,%<br>(10/10) | 0.89 ± 0.10*,@,\$,%<br>(10/10) | 0.85 ± 0.11*,@,%<br>(10/10) | 3.42 ± 0.33 |

Abbreviations: Dap, daptomycin
\*p-value <0.05 LSVT-1701 treatment groups compared to untreated control; @p-value <0.05 LSVT-1701 treatment groups compared to daptomycin alone; #p-value <0.05 LSVT-1701 32.5 mg/kg iv QD for 4 d compared to iv QD for 2 d and single dose; \$p-value <0.05 LSVT-1701 50 mg/kg iv QD for 4 d compared to single dose; %p-value <0.05 LSVT-1701 50 mg/kg iv QD for 4d compared to 32.5 mg/kg iv QD for 2 d and single dose.

The mean C_max_ (μg/mL), AUC_0-last_ (μg.h/mL), and t_1/2_ (h) for the LSVT-1701 32.5 mg/kg group were 14.5, 12.1, and 0.525, respectively, and for the 50 mg/kg group were 24.0, 20.8, and 0.555, respectively.

## Discussion

In this MRSA IE model, both LSVT-1701 doses tested, when administered daily as a 1-hour iv infusion, for 4 d in combination with daptomycin, microbiologically sterilized all target organs tested. This is particularly impressive, given the high bio-burdens in such target tissues at the time of therapy initiation (i.e., > 8 log_10_ CFU/g in vegetations and > 6 log_10_ CFU/g in kidneys and spleen).

Prior studies determined that the AUC/MIC ratio is the strongest predictor of efficacy of an antistaphyloccal lysin in a *S. aureus* mouse thigh infection model; in addition, time-above-MIC (T>MIC) was also predictive of outcome in this same model (10). In the current study, the mean AUC values for 50 mg/kg and 32.5 mg/kg dose-regimens given as four daily doses were 20.8 and 12.1 μg.h/ml, respectively. Based on a Phase 1 study in healthy volunteers with LSVT-1701, single 6 and 10 mg/kg doses administered iv over 1 h resulted in mean AUC_0-last_ values of 24.6 and 59.8 μg.h/ml, respectively; exceeding the apparent pharmacologically relevant “efficacious exposure” (which resulted in sterility of all target tissues at 32.5 mg/kg) by 2-5-fold in the IE model. In addition, the AUC exposures of a single 6 mg/kg dose of LSVT-1701 is within the exposure margins relative to the ‘no observed adverse effect level’ (NOAEL) dose level in a 28-day GLP toxicology study in nonhuman primates (*unpublished data*). Furthermore, LSVT-1701 can be dosed for multiple days to maximize efficacy potential in alignment with the posology used in this rabbit IE model.

Of note, the same rabbit IE model caused by the same MRSA strain (MW2) as used in the present investigation was recently employed to study another anti-staphylococcal phage lysin, CF-301 (Exebacase^®^) [8]; this lysin is currently in a Phase 3 clinical trial for the treatment of MRSA bacteremia including right-sided IE (clinicaltrials.gov identifier NCT04160468). Despite utilizing the identical daptomycin dose-regimen as in the current study (4 mg/kg iv QD for 4 d), the same high rates of target tissue sterilization were not observed in the CF-301 studies [8]. Although both CF-301 and LSVT-1701 are recombinant anti-staphylococcal lysins, there are distinct structural differences. CF-301 is 26 kDa, has only one catalytic domain (an endopeptidase), and is limited to a single-dose regimen based on preclinical toxicology studies (dose-limiting toxicity of vasculitis and hypersensitivity) (11). In contrast, LSVT-1701 is 54 kDa, has two catalytic domains (an endopeptidase and an amidase) which cleave at two sites in the cell wall; it can also be administered as multiple doses, with no similar dose-limiting toxicities observed in a 4-week GLP repeat-dose study in rats and non-human primates) (12,13). Because of these differences, LSVT-1701 will be evaluated in an upcoming Phase 2b study of *S. aureus* bacteremia, including both right-sided and left-sided IE.

There are several limitations to the current study. *First*, only one MRSA strain was assessed. Future studies will require multiple strains of both MRSA and MSSA to be evaluated in this same model. *Second*, although not the objective of this study, the rates and extents of microbiologic target tissue relapse following the 4 d daptomycin + LSVT-1701 regimens were not evaluated. Such investigations of relapse are planned for future studies. *Third*, while a hybrid immunoaffinity-LC-MS/MS method was used to quantify LSVT-1701 in rabbit plasma, the Phase 1 PK determinations were made with an ELISA-based method. Subsequent clinical PK evaluations will be performed with an immunoaffinity-LC-MS/MS method.

In conclusion, LSVT-1701 administered at 32.5 or 50 mg/kg in a 4 d daily-dose regimen in combination with daptomycin resulted in microbiologic sterilization of all target organs in this MRSA IE model. These data support further clinical development of LSVT-1701 for the treatment of MRSA endovascular infections including IE.

## Acknowledgements

We are grateful for the help of Surya Chitra, a consultant for Lysovant, for statistical support. These studies were supported by a research grant to ASB from Lysovant.

## Materials and Methods

### Bacterial strain

MRSA strain MW2 (USA 400; clinical complex [CC] 1) was used because this isolate: is clinically-derived; genome-sequenced; represents a common hospital-acquired MRSA clonotype; is virulent in experimental IE models; and is daptomycin-susceptible *in vitro* (14,15). Moreover, the dose of daptomycin that yields substantial, but moderate clearance of MW2 from target tissues, has been well-established in prior studies (8,9). This provides a key microbiologic ‘window’ to disclose the synergistic potential of adjunctive agents in this model. The daptomycin MIC of MW2, as determined by standard CLSI broth dilution methods, is 1 μg/mL, and that of LSVT-1701 is 0.5 μg/ml as determined by broth microdilution in cation-adjusted (CA)-MHB + 0.25mM dithiothreitol (DTT) at an incubation time of 20 h with visual and/or spectrophotometric endpoints to assess MIC. These conditions are in line with the current CLSI M07 guidelines which recommend the use of CA-MHB and incubation for 20 h for susceptibility testing of *S. aureus*. The addition of 0.25 mM DTT decreases assay variability.

#### In vivo efficacy of LSVT-1701 in the rabbit model of IE

The standard MRSA aortic valve IE rabbit model was used to compare *in vivo* efficacy profiles of two-dose regimens of LSVT-1701 (32.5 mg/kg and 50 mg/kg), given by varying dose schedules, in combination with daptomycin (vs daptomycin alone). Briefly, an indwelling transcarotid artery-to-left ventricle catheter was inserted under general anesthesia with xylazine and ketamine to induce sterile aortic vegetations. At 48 h after catheter placement, IE was induced by an iv challenge of ~ 2 x 10^5^ cfu, an inoculum, which induces IE in >95% of catheterized rabbits with this strain based on extensive prior data (8, 9, 16–18). One group of untreated animals (controls) were humanely sacrificed by iv pentobarbital at ~24 h post-infection, target organs sterilely removed and quantitatively cultured to establish the baseline MRSA bioburden in these tissues (vegetations; kidneys; and spleen). In the remaining treatment animals, similar sacrifices were done, and quantitative cultures performed as above at 24 h after the last dose of daptomycin and/or LSVT-1701 to minimize any antimicrobial carryover impacts on cultures. In addition, baseline quantitative blood cultures were done post-infection, as well as post-therapy. Microbiologic data in the various treatment groups for each target organ was calculated as either the mean log_10_ CFU/ml (+/-standard deviation [SD]; for blood) or the mean log_10_ CFU/g of tissue (+/-SD; for target tissues). Experimental IE studies were approved by the IACUC Committee of The Lundquist Institute prior to performance.

#### Treatment regimens of LSVT-1701 +/- daptomycin

At 24 h after induction of IE, 10 animals each were randomized into one of the seven treatment groups: **i**) untreated controls (sacrificed at 24 h post-infection); **ii**) daptomycin at 4 mg/kg iv once daily (QD) by bolus x 4 d; **iii**) LSVT-1701 at 32.5 mg/kg iv given as a single dose + daptomycin 4 mg/kg iv QD x 4 d; **iv**) LSVT-1701 at 50 mg iv single dose + daptomycin 4 mg/kg iv QD x 4 d; **v**) LSVT-1701 at 32.5 mg/kg QD x 2 d + daptomycin 4mg/kg iv QD x 4 d; **vi**) LSVT-1701 at 32.5 mg/kg QD x 4 d + daptomycin 4 mg/kg iv QD x 4 d; and **vii**) LSVT-1701 at 50 mg/kg iv QD x 4 d + daptomycin 4 mg/kg iv QD x 4 d. The LSVT-1701 iv dose was given by an iv infusion of 1 h duration administered within 30 min following the AM daptomycin iv bolus dose. We did not include a LSVT-1701-alone group, as pilot IE studies found no microbiologic impacts with this regimen (*data not shown*).

#### LSVT-1701 pharmacokinetic and pharmacodynamics

A PK study was conducted in rabbits with the same IE model, with LSVT-1701 administered at 32.5 and 50 mg/kg via 1-hour iv infusion. Serum PK samples (n=3/timepoint/group) were collected at 1, 3, 15, 30, 60, 180, and 360 min following the conclusion of the infusion interval.

LSVT-1701 was quantified in rabbit serum samples using an affinity purification mass-spectrometry (AP-MS) method. Briefly, LSVT-1701 was immunoprecipitated from serum samples using a biotinylated mouse anti-LSVT-1701 monoclonal antibody in conjunction with streptavidin-coated beads. Following overnight trypsin digestion of LSVT-1701, proteolytic fragments of the lysin were subjected to LC-MS/MS analysis and test article concentrations were determined via interpolation from calibration standards. Pharmacokinetic parameters were calculated with Phoenix WinNonlin v 8.3.1 from the resultant concentration-time data points. Pharmacokinetic data presented here were calculated from the stand-alone rabbit PK study, as these data offered greater sampling density and provided earlier sample collection time points than those available during the conduct of the IE model.

#### Statistical analyses

These data (primary end-points) were statistically compared by a mixed effect model treatment groups as fixed effects, and rabbits nested within each treatment as random effect (using SAS^®^ version 9.4 or later). The *P* values were adjusted for multiple comparisons using Tukey-Kramer method. *P* values of < 0.05 were considered statistically significant.

## References

1. Lowy FD. *Staphylococcus aureus* infections. 1998. N Engl J Med 339:520–32. doi: 10.1056/NEJM199808203390806.

2. Tong SY, Davis JS, Eichenberger E, Holland TL, Fowler VG Jr. 2015. *Staphylococcus aureus* infections: epidemiology, pathophysiology, clinical manifestations, and management. Clin Microbiol Rev 28:603–61. doi: 10.1128/CMR.00134-14.

3. Holland TL, Baddour LM, Bayer AS, Hoen B, Miro JM, Fowler VG Jr. 2016. Infective endocarditis. Nat Rev Dis Primers 2:16059. doi: 10.1038/nrdp.2016.59.

4. Sakoulas G, Moise-Broder PA, Schentag J, Forrest A, Moellering RC Jr, Eliopoulos GM. 2004. Relationship of MIC and bactericidal activity to efficacy of vancomycin for treatment of methicillin-resistant *Staphylococcus aureus* bacteremia. J Clin Microbiol 42:2398–2402.

5. Soo Youn Jun, Gi Mo Jung, Seong Jun Yoon, Myoung-Don Oh, Yun-Jaie Choi, Woo Jong Lee, Joon-Chan Kong, Jae Goo Seol, Sang Hyeon Kang. 2013. Antibacterial properties of a preformulated recombinant phage endolysin, SAL-1. Int J Antimicrob Agents 41:156–61. doi: 10.1016/j.ijantimicag.2012.10.011.

6. Kim NH, Park WB, Cho JE, Choi YJ, Choi SJ, Jun SY, Kang CK, Song KH, Choe PG, Bang JH, Kim ES, Park SW, Kim NJ, Oh MD, Kim HB. 2018. Effects of phage endolysin SAL200 combined with antibiotics on *Staphylococcus aureus* infection. Antimicrob Agents Chemother 62:e00731–18. doi: 10.1128/AAC.00731-18.

7. Fowler VG Jr, Boucher HW, Corey GR, Abrutyn E, Karchmer AW, Rupp ME, Levine DP, Chambers HF, Tally FP, Vigliani GA, Cabell CH, Link AS, DeMeyer I, Filler SG, Zervos M, Cook P, Parsonnet J, Bernstein JM, Price CS, Forrest GN, Fätkenheuer G, Gareca M, Rehm SJ, Brodt HR, Tice A, Cosgrove SE; S. aureus Endocarditis and Bacteremia Study Group. 2006. Daptomycin versus standard therapy for bacteremia and endocarditis caused by *Staphylococcus aureus*. N Engl J Med 355:653–65. doi: 10.1056/NEJMoa053783.

8. Indiani C, Sauve K, Raz A, Abdelhady W, Xiong YQ, Cassino C, Bayer AS, Schuch R. 2019. The antistaphylococcal lysin, CF-301, activates key host factors in human blood to potentiate methicillin-resistant *Staphylococcus aureus* bacteriolysis. Antimicrob Agents Chemother 63:e02291–18. doi: 10.1128/AAC.02291-18.

9. Shah SU, Xiong YQ, Abdelhady W, Iwaz J, Pak Y, Schuch R, Cassino C, Lehoux D, Bayer AS. 2020. Effect of the lysin Exebacase on cardiac vegetation progression in a rabbit model of methicillin-resistant Staphylococcus aureus endocarditis as determined by echocardiography. Antimicrob Agents Chemother 64:e00482–20. doi: 10.1128/AAC.00482-20.

10. Rotolo JA, Ramirez RA, Schuch R, Machacek M, Khariton T, Ghahramani P, Wittekind M. 2020. PK-PD driver of efficacy for CF-301, a novel anti-staphylococcal lysin: implications for human target dose. ASM.

11. https://www.fda.gov/media/115771/download

12. Jun SY, Jung GM, Yoon SJ, Choi YJ, Koh WS, Moon KS, Kang SH. 2014. Preclinical safety evaluation of intravenously administered SAL200 containing the recombinant phage endolysin SAL-1 as a pharmaceutical ingredient. Antimicrob Agents Chemother 58:2084–8. doi: 10.1128/AAC.02232-13.

13. Jun SY, Jung GM, Yoon SJ, Youm SY, Han HY, Lee JH, Kang SH. 2016. Pharmacokinetics of the phage endolysin-based candidate drug SAL200 in monkeys and its appropriate intravenous dosing period. Clin Exp Pharmacol Physiol 43:1013–6. doi: 10.1111/1440-1681.12613.

14. Mishra NN, Rubio A, Nast CC, Bayer AS. 2012. Differential adaptations of methicillin-resistant Staphylococcus aureus to serial in vitro passage in daptomycin: evolution of daptomycin resistance and role of membrane carotenoid content and fluidity. Int J Microbiol 2012:683450. doi: 10.1155/2012/683450.

15. Mishra NN, Yang SJ, Sawa A, Rubio A, Nast CC, Yeaman MR, Bayer AS. 2009. Analysis of cell membrane characteristics of in vitro-selected daptomycin-resistant strains of methicillin-resistant *Staphylococcus aureus*. Antimicrob Agents Chemother 53:2312–8. doi: 10.1128/AAC.01682-08.

16. Bayer AS, Lam K. Bayer AS. 1985. Efficacy of vancomycin plus rifampin in experimental aortic-valve endocarditis due to methicillin-resistant *Staphylococcus aureus*: in vitro-in vivo correlations. J Infect Dis 151:157–65. doi: 10.1093/infdis/151.1.157.

17. Bayer AS, Greenberg DP, Yih J. 1988. Correlates of therapeutic efficacy in experimental methicillin-resistant Staphylococcus aureus endocarditis. Chemotherapy 34:46–55. doi: 10.1159/000238547.

18. Trotonda MP, Xiong YQ, Memmi G, Bayer AS, Cheung AL. 2009. Role of mgrA and sarA in methicillin-resistant Staphylococcus aureus autolysis and resistance to cell wall-active antibiotics. J Infect Dis 199:209–18. doi: 10.1086/595740.

